# Acute and chronic stage adaptations of vascular architecture and cerebral blood flow in a mouse model of TBI

**DOI:** 10.1101/479626

**Authors:** Joe Steinman, Lindsay S. Cahill, Margaret M. Koletar, Bojana Stefanovic, John G. Sled

## Abstract

The 3D organization of cerebral blood vessels determines the overall capacity of the cerebral circulation to meet the metabolic requirements of the brain. This study used Arterial Spin Labeling (ASL) MRI with a hypercapnic challenge and ex vivo Serial Two-Photon Tomography (STPT) to examine the relationship between blood flow and 3D microvascular structure following traumatic brain injury (TBI) in a mouse. Mice were exposed to a controlled cortical impact TBI and allowed to recover for either 1 day or 4 weeks. At each time point, ASL MRI was performed to quantify cerebral perfusion and the brain vasculature was imaged in 3D with STPT. Registration of ASL to STPT enabled flow changes to be related to the underlying microvascular structure in each ASL voxel. Hypoperfusion under rest and hypercapnia was observed both 1 day and 4 weeks post-TBI. Vessel density and vascular volume were reduced 1 day post-TBI, recovering by 4 weeks; however, the reorganized vasculature at the latter time point possessed an abnormal radial pattern. Our findings demonstrate functionally significant long-term changes in the vascular architecture following injury and illustrate why metrics beyond traditional measures of vessel density are required to understand the impact of vascular structure on function.

## Introduction

The 3D organization of vessels is an important factor in determining cerebral blood flow (CBF), with the vessel connections, shape, length, and diameter determining overall network resistance [1]. While regions of higher metabolic activity often show increased vessel density [2], substantial increases in density may not elevate flow by an equivalent amount [3]. This suggests that to understand the relationship between CBF and vascular structure, quantitative metrics beyond 2D measures of vessel density are needed. This vascular structure-blood flow relationship is thought to be important in a number of brain pathologies, including traumatic brain injury (TBI), where the vasculature undergoes significant damage, vessel loss, and regrowth [4, 5, 6, 7, 8].

Past studies have correlated CBF changes or functional outcome with vascular structural changes in TBI [9, 10], focusing on histological measures of density and diameter. Such 2D metrics of vascular morphology do not always correlate with blood flow. For example, vessel density in a hippocampal region correlated with CBF early post-injury in a lateral fluid percussion injury in rats [5]. At 8 months in the same model, there was increased vessel density in perilesional cortex and reduced flow, with bilateral decreases in hippocampal CBF occurring without changes in vessel number [9].

Quantification of blood flow and vascular architecture is limited by the lack of availability of high-resolution 3D imaging techniques that may be combined with in vivo functional imaging. While optical clearing may be used to enhance light penetration, solvents often distort tissue and cause optical aberrations. A methodology with the ability to image the microvasculature in 3D over a large volume of tissue without optical clearing would overcome these deficiencies.

Automated techniques have been developed which image large regions of the murine brain without optical clearing [11]. One such technique, Serial 2-Photon Tomography (STPT), incorporates a 2-photon fluorescence microscope and vibratome [12]. The vibratome slices through tissue, enabling deep tissue imaging. To date, most STPT studies have acquired images at cellular-resolution in-plane with low through-plane resolution, typically with 75 – 100 µm spacing between coronal sections. This method enables whole-brain imaging in under 12 hours [13], but reduced through-plane resolution is not amenable to 3D vascular analysis.

In the present study, we used a combination of 2D Arterial Spin Labeling (ASL) MRI and STPT to measure changes in cerebrovascular function and to characterize the 3D vascular architecture in adult mice post-TBI. The standard STPT protocol was modified to obtain images at an isotropic voxel resolution of 2 µm, enabling 3D visualization and quantification of the geometry and branching patterns of the vascular network. The differences in vascular structure were then correlated with the functional measures of flow and vascular reserve. These techniques highlight unique features of vascular architecture and blood flow, providing improved understanding of the role of the microvasculature in TBI recovery.

## Materials and Methods

### Animals

A total of 40 mice (19 male, 21 female, 12 - 13 weeks old, mean weight 24.2 ± 0.6 g), Cre x tdTomato (B6.Cg-Tg(Tek-cre)1Ywa/JxB6; 129S6-Gt(ROSA)26Sortm14(CAG-tdTomato)Hze/J), bred in-house, were used [14]. See Supplementary Table 1 for a detailed description of animal allocation within the study. The mice express a variant of red fluorescent protein in endothelial cells, under the direction of the Tie2 promoter [15]. Mice received either a controlled cortical impact (CCI) or craniotomy without impact (sham). They were permitted to recover 24 hours or 4 weeks post-injury. Mice were randomly assigned into one of four groups: TBI-1-Day, TBI-4-Weeks, Sham-1-Day, and Sham-4-Weeks. All animal experiments were approved by the Animal Care Committee at the Toronto Centre for Phenogenomics. Animal experiments were conducted in accordance with the Canadian Council on Animal Care’s guide to the Care and Use of Experimental Animals and complied with the ARRIVE guidelines.

### Controlled Cortical Impact

Mice were anaesthetized with isoflurane (5 % induction, 1.5 - 2 % maintenance) in 100 % O_2_. They were administered 5 mg/kg Baytril (Rompun, Bayer Inc., Toronto, Canada) and 1.2 mg/kg slow release buprenorphine (Chiron Compounding Pharmacy Inc., Guelph, Canada) subcutaneously, in addition to 1.5 mL of subcutaneous sterile saline (0.9 % NaCl). Mice were stabilized in a stereotaxic frame. Body temperature was maintained via a heating pad set to 37 °C. Immediately prior to scalp incision, mice were administered 0.2 mL of 1 % lidocaine (AstraZeneca, Mississauga, Canada) subcutaneously over the skull. The scalp was shaved, then cleaned with alcohol and betadine solutions. A craniotomy approximately 2.2 mm in diameter was performed and centred at 1.5 mm posterior to bregma and 1.7 mm lateral to the midline on the left hemisphere. A TBI (1.5 mm diameter tip, 1 mm depth, 2 m/s speed, 200 ms dwell time) was delivered with an electromagnetically driven piston [16]. The craniotomy was sealed with a 5 mm diameter glass coverslip (World Precision Instruments, Sarasota, Florida, USA). The scalp was sutured; betadine and polysporine were applied to the incision. Each mouse was individually housed in a sterile cage with DietGel (76 A, PharmaServ, Framingham, MA, USA) and food pellets placed at the bottom of the cage. Mice had ad libitum access to food and water in a pathogen-free environment on a 12-hour light: dark cycle. Identical procedures were performed for shams, with the exception of the impact.

### Measurement of CBF with continuous Arterial Spin Labeling (CASL)

Prior to CASL imaging, in vivo ultrasound imaging under isoflurane was performed as described previously [17]. The ultrasound data is beyond the scope of the current study and will be the subject of future investigations.

The ASL procedure followed that described in Cahill et al. [17] and is summarized below.

#### Imaging preparation and hypercapnia protocol

Immediately following ultrasound, mice were endotracheally intubated and mechanically ventilated under 1 % isoflurane in 100 % O_2_. 0.1 mg/kg pancuronium bromide (Sigma-Aldrich, St. Louis, USA) was injected intraperitoneally for muscle relaxation and optimal breathing control during imaging. Partial pressure of carbon dioxide, respiration rate, and temperature were monitored. Each mouse underwent a CO_2_ challenge, composed of an inhaled gas mixture alternating between rest (30 % O_2_ : 70 % N_2_) and hypercapnia (5 % CO_2_ : 30 % O_2_ : 65 % N_2_). In total, there were 5 – 6 scans per mouse, with two of the scans under hypercapnia. Following hypercapnia, there was a 6-minute break between measurements to allow for return to normal physiological state.

#### ASL

2D CASL measurements were performed on a multi-channel 7 T scanner (Magnex Scientific, Oxford, UK). A coronal slice intercepted the centre of the TBI site and the labeling plane was positioned posterior to the imaging slice by approximately 0.8 cm. Each repetition included a 3-s 9-µT RF labelling pulse and a 500 ms post-label delay. The CASL images were acquired using a 2D fast spin-echo sequence with TR = 6000 ms, TE_eff_ = 15ms, echo train length = 16, slice thickness = 2 mm, matrix size 200 × 384, in-plane resolution of 250 µm. CBF was calculated with a single-compartment biophysical model with correction for the magnetization transfer enhancement of T_1_ relaxation [17, 18]. As TBI may cause edema [19] and influence tissue T_1_, affecting CBF measurements [18], T_1_ was mapped in the same 2 mm-thick coronal section using an inversion recovery fast spin-echo sequence (TR = 5000 ms, TE = 11ms, inversion times = 50, 100, 200, 400, 800, 1600, 3200 ms).

### T_2_-weighted whole brain in vivo anatomical imaging

T_2_-weighted whole brain images were acquired using a 3D fast spin-echo sequence with the following parameters: TR = 1800 ms, TE_eff_ = 40 ms, echo train length = 12, matrix size 120 × 144 × 72, isotropic image resolution = 292 µm. The acquired image was used as an intermediary in a registration pipeline to align the STPT images to ASL data (see below).

### Brain sample preparation

The perfusion protocol have been described elsewhere [20, 21, 22] and is briefly summarized here. Immediately following in vivo MRI, mice were anaesthetized with an IP injection of 100 µg/g ketamine and 20 µg/g xylazine. The following solutions were perfused through the left ventricle, with 2 mM ProHance (Bracco Diagnostics Inc., Monroe Plains, New Jersey, USA) added to each solution for MR contrast: 30 mL of 0.1 M PBS (Wisent Inc., Quebec, Canada) containing 1 µL/mL heparin (1000 USP units/mL); 20 mL of 4 % paraformaldehyde (PFA) (Electron Microscopy Sciences, Hatfield, Pennsylvania, USA); and 20 mL of a 2 % gelatin solution (Sigma, St. Louis, USA) containing 0.5 % (w/v) FITC-conjugated albumin (Sigma) [22]. Mice were decapitated and the brains within the skull were fixed overnight at 4° C in 4 % PFA with 2 mM ProHance. Brains were transferred to PBS containing 2 mM ProHance and 0.02 % sodium azide for at least one week [23].

### Ex vivo whole-brain MRI

Brains were scanned in a 16-coil solenoid array [24] using a T_2_-weighted 3D fast spin-echo sequence with a cylindrical *k*-space acquisition [25]. Sequence parameters were 4 averages, TR = 350 ms, TE_eff_ = 30 ms, echo train length = 6, matrix size 504 × 504 × 630, isotropic image resolution = 40 µm.

### 3D Serial Two-Photon Tomography

STPT was imaged on two channels as previously described [13]. Light emitted from the specimen was split using a 516 nm single-edge dichroic beamsplitter (Semrock FF516-di01-25 × 36). Due to the greater signal strength, only the data from the channel with light greater than 516 nm wavelength were analyzed: this channel provided signal from both the tomato fluorescent protein and the gelatin: FITC-conjugate.

2-photon fluorescence imaging commenced 50 µm below the cut surface and involved collecting a mosaic of overlapping tiles. Each field-of-view (FOV) was sampled with 832 voxels along × and y for a 1.37 µm in-plane imaging resolution and acquired through a tissue depth of 100 µm, with 2 µm separation between tiles. Following acquisition of all 3D images within the xy-plane, the tissue was sliced 30 µm by a vibratome integrated with the translation stage and the sample moved towards the objective. A typical imaging session involved 150 vibratome sections, corresponding to a tissue volume of 4 mm (*x*) × 4 mm (*y*) × 4.5 mm (*z*) and an imaging time of approximately 33 hours.

### STPT data processing and reconstruction

Each 3D image was corrected for uneven illumination using an agar phantom loaded with FITC. The 3D images were blurred in the xy-plane by a Gaussian with full-width-half-maximum (FWHM) of 2 µm and resampled to an isotropic voxel size of 2 × 2 × 2 µm^3^. Image stitching was performed with a Fiji/ImageJ plug-in based on the algorithm of Preibisch et al. [26], with overlapping 3D images merged via a linear blending algorithm.

### Image Registration Pipeline

Registration of the ASL data to the STPT images was performed using in vivo and ex vivo whole-brain structural MR images as intermediate targets for registration as illustrated in Figure 1.

**Figure 1.**
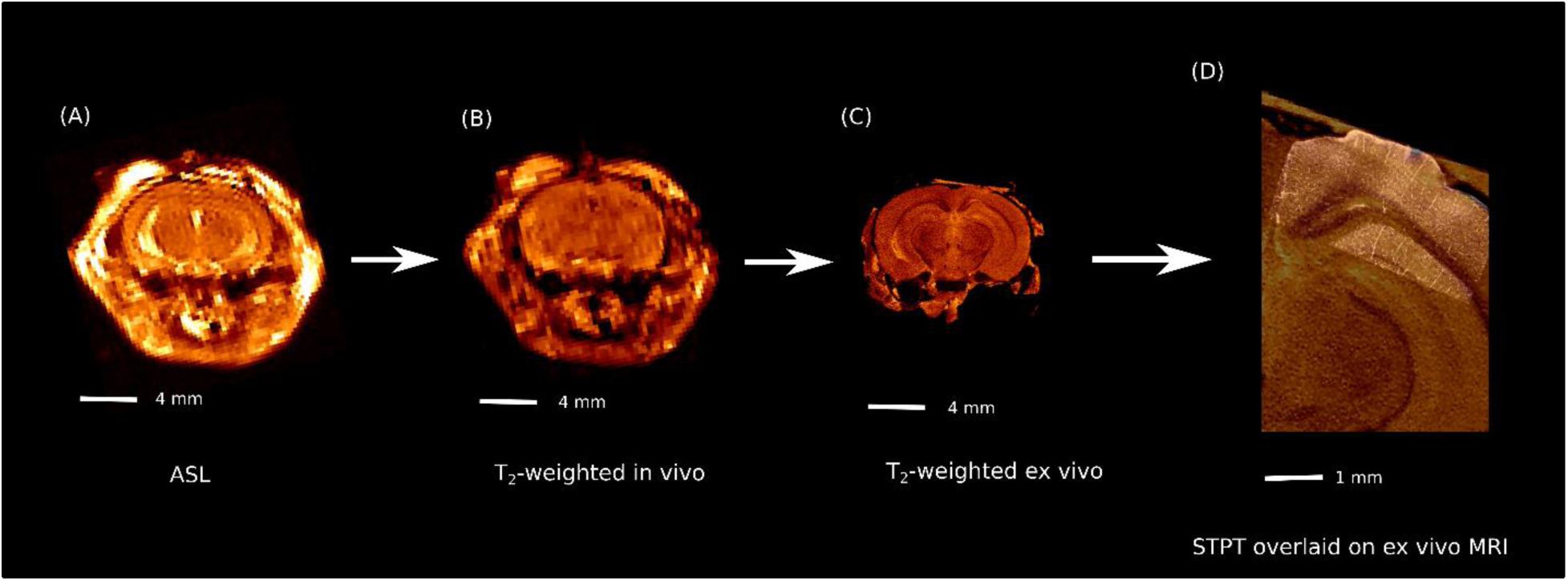
(A) Representative T_2_-weighted anatomical image from the ASL acquisition (B) Single-slice from in vivo T_2_-weighted whole-brain image (C) Single-slice from ex vivo T_2_-weighted whole-brain with the skin and overlying tissue from the skull removed (D) Single-slice STPT, showing the ipsilateral cortex and part of the hippocampus overlaid on the ex vivo MR. To register the CBF maps to the stitched STPT images, the in vivo and ex vivo whole-brain structural MR images were used as intermediates in a registration pipeline based on manual identification of common landmarks between the respective images.

### Vessel tracking

The vessel tracking segmentation algorithm has been described elsewhere [20, 27]. Tracking was initiated through automated placement of multiple seeds inside each putative vessel [14, 20]. Prior to tracking, cortical voxels in the ASL data were manually labelled by overlaying the control ASL image onto the STPT data. Cortical voxels were subdivided into two groups: (A) those located within the craniotomy region (TBI site) through the cortical depth, termed the ‘ipsilateral’ injury region; and (B) those outside the TBI site, termed the ‘perilesional’ region. Ipsilateral voxels that corresponded to the centerline of the injury were further labelled ‘centerline’. The centerline voxels were later used to calculate the distance of the cortical voxels from the injury center (see below). Each voxel in the STPT image was assigned a label of ipsilateral or perilesional, based on the cortical ASL voxel type to which it corresponded. Only these voxels were tracked.

The datasets were pruned during post-processing to remove short, false vessels (‘hairs’) which are either disconnected from the vessel network or connected at only a single end to a parent vessel. These vessels are produced either by noise at a vessel boundary that is interpreted as a separate vessel or by the anisotropy of the 2-photon fluorescence data causing short vessels to be traced parallel to the optical axis.

### Data analysis

#### Quantification of vessel network properties

Vessel diameters calculated were the outer diameter, which is the sum of the lumen (inner) diameter and the vessel wall thickness. Vessels with diameters less than 8 µm were labelled capillaries, since capillary cut-offs in the literature typically range from 6 µm to 10 µm [14, 22]. All vessels traced emitted fluorescence greater than 516 nm and were either tomato-fluorescent and gel-perfused, or only tomato-fluorescent in the event that a given vessel was not perfused.

‘Centerline voxels’ in the ASL data (defined in *Vessel tracking*) were defined as 0 mm from the injury centre. The distance of all other cortical ASL voxels was defined by the Euclidean distance to the nearest centerline voxel. Each vessel segment in the SPTP data was assigned a nearest ASL voxel and distance from injury centre based on the position of the vessel segment’s midpoint. The vessel length density at each voxel was the sum of the lengths of all vessels within an ASL voxel divided by the voxel volume.

#### Quantification of radial vascular architecture

The vascular network at 4 weeks post-TBI (the TBI-4-Weeks group) possessed a different patterning relative to the other groups (TBI-1-Day, Sham-1-Day, Sham-4-Weeks). From the center of the injury, vessels emanated outwards in a radial pattern. To quantify this feature, a marker was placed at the centre of the craniotomy on the cortical surface. Each segmented vessel was considered as a straight line. Another line was defined connecting the vessel midpoint and marker. The angle between these two lines was calculated for all vessels, averaged, and compared between mice and groups. This metric is termed here as the extent of *radialness.* Vessel networks that display a radial pattern would be expected to possess a smaller mean angle.

#### Statistical analysis

All data are reported as mean ± standard error. Data analysis was conducted in Python and R. To compare mean differences between groups, the dependent variable was modelled as a linear function of group. An ANOVA was performed on the linear model with p < 0.05 interpreted as significant. If p < 0.05, a Tukey Honestly Significant Differences post-hoc test was conducted to assess between group differences.

To assess if TBI influenced CBF, an ANOVA was performed on a linear mixed effects model where CBF was modeled as a three-way interaction between CO_2_ state (rest vs. hypercapnia), time (1 day vs. 4 weeks), and treatment (TBI vs. Sham), with specimen as the random factor to account for repeated measures. For analysis of CBF as a function of distance from the injury centre, CBF vs. distance plots for each group were fit with a cubic spline with 3 degrees of freedom. An ANOVA was performed on a linear mixed effects model where CBF was modeled as a four-way interaction between distance (which was fit with a cubic spline), state (rest vs. hypercapnia), time (1 day vs. 4 weeks), and treatment (TBI vs. Sham), with specimen as the random factor. Coefficient of variation (COV) was defined as the ratio between the standard deviation of CBF in ASL cortical voxels divided by their mean value, times 100 %. An ANOVA was performed on a linear mixed effects model where COV was modeled as a four-way interaction between distance, CO_2_ state, time, and treatment, with specimen as the random factor. For analysis of mean vessel length density, microvascular volume, extravascular distance, capillary diameter, and radialness within the TBI site, an ANOVA was performed on a linear model where these variables were modeled as a two-way interaction between time and treatment. To assess how density, volume, and extravascular distance changed as a function of distance from the injury centre, an ANOVA was performed on a linear mixed effects model where these variables were modeled as a three-way interaction between distance, time, and treatment.

## Results

### STPT images of the microvasculature demonstrate abnormal morphology following TBI

TBI-1-Day mice showed loss of vessels and altered vessel morphology (Figure 2). Panel 2A is a maximum intensity projection (MIP) of a coronal section of the cerebral cortex in a mouse that did not undergo craniotomy. Note the uniform vessel density and regular pattern of the vessel network. This contrasts with Panel 2C (MIP of a TBI-1-Day mouse), where fewer vessels are detected inside the injury core. Outside the core, many penetrating vessels are highly tortuous (Panel 2B).

**Figure 2.**
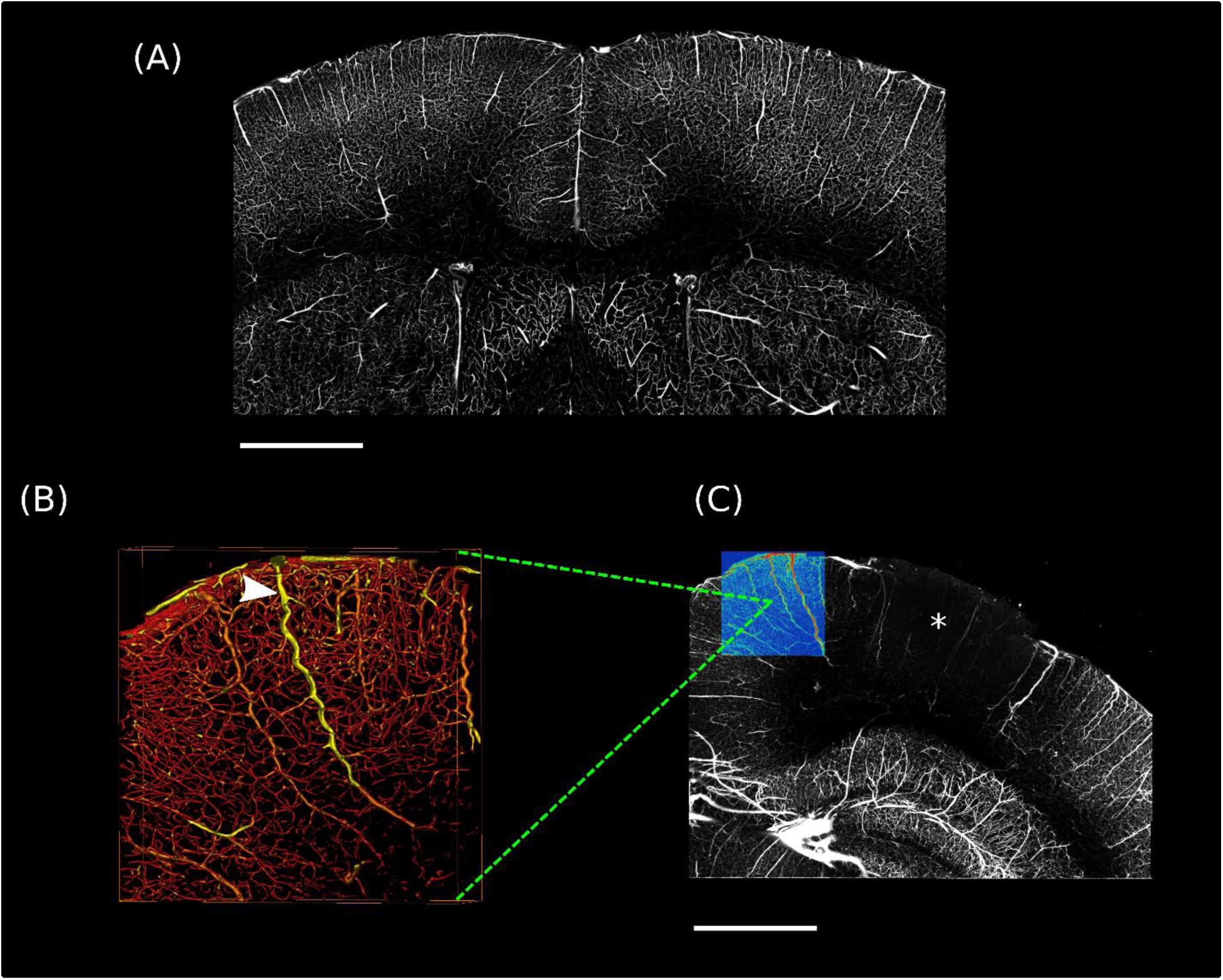
(A) Maximum Intensity Projection (MIP) through 100 µm of tissue, acquired from a mouse that did not undergo craniotomy. Scale bar = 1 mm (B) and (C) Data acquired from a TBI-1-Day mouse. (B) is an isosurface rendering of 200 µm tissue thickness corresponding to the colored region in (C). (C) is a MIP through 400 µm of tissue. Note the presence of tortuous (twisted) vessels in regions surrounding the impact core (arrowhead in B) and the reduction in vessel density within the core (asterisk in C). Scale bar = 1 mm. All data acquired with STPT.

### Hypoperfusion in TBI persist up to 4 weeks

Figures 3A and 3B plot the mean CBF in the ipsilateral cortex (craniotomy region) under rest and hypercapnia. CBF depended significantly on CO_2_ state (F = 87.5, p = 8.4 × 10^−11^) and treatment (F = 15.1, p = 0.00047 respectively), but not on time (F = 0.3, p = 0.60). There was a significant interaction between state and time (F = 6.1, p = 0.019).

**Figure 3.**
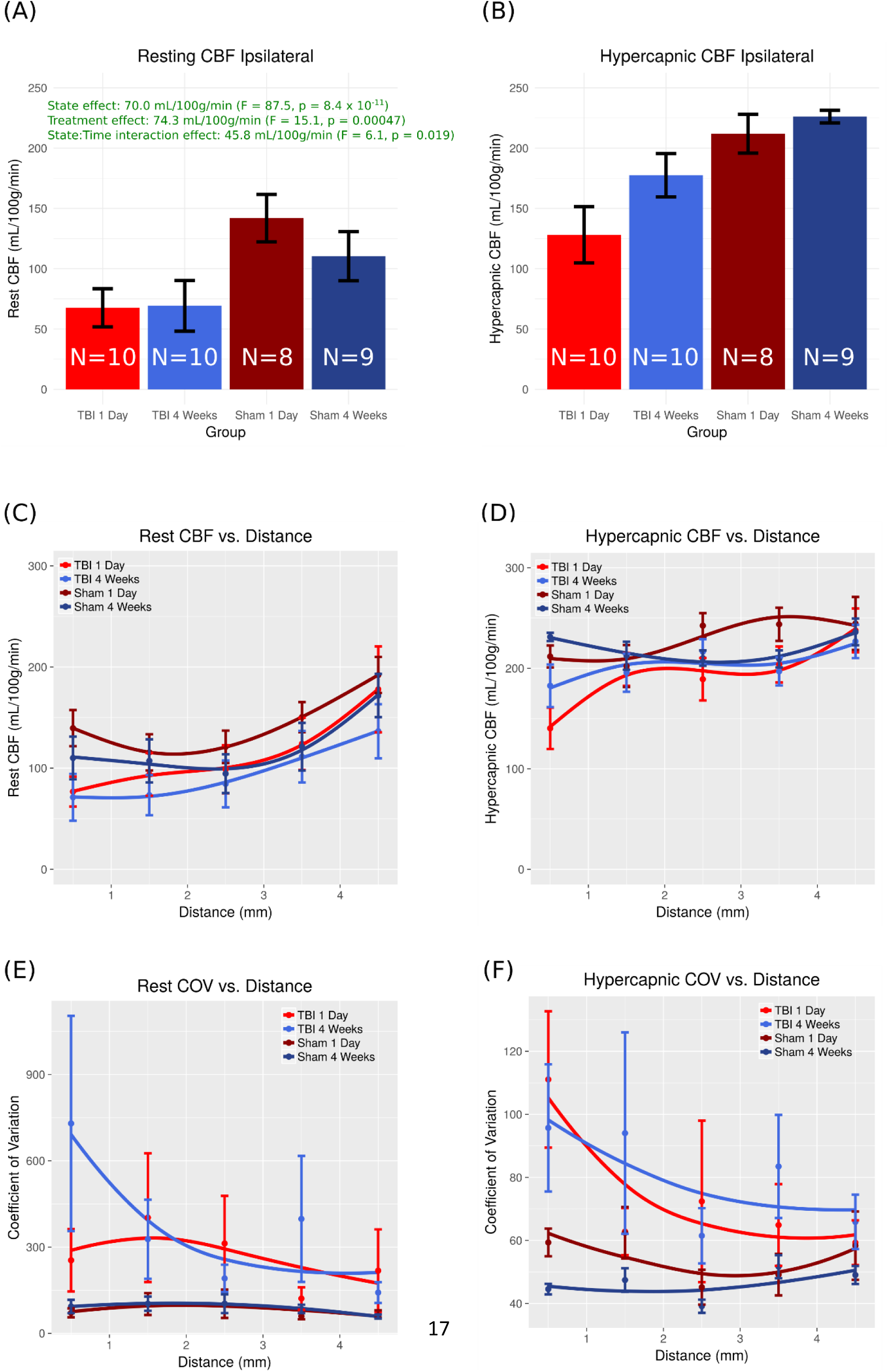
Mean CBF calculated with ASL MRI in the ipsilateral cortex under (A) rest and (B) hypercapnia. CBF as a function of distance from the injury centre on the injured side under (C) rest and (D) hypercapnia. Coefficient of Variation as a function of distance from the injury centre under (E) rest and (F) hypercapnia. Data points are the mean ± SEM, and fitting was performed with Loess regression. N refers to the number of mice.

To investigate CBF outside the injured core, Figures 3C and 3D plot the CBF over the entire injured cortex as a function of distance from the injury centre under rest and hypercapnia respectively. There was a significant dependence on distance (F = 30.4, p < 2.2 × 10^−16^), state (F = 89.8, p < 2.2 × 10^−16^), and treatment (F = 9.4, p = 0.0033), but no time-dependence (F = 0.12, p = 0.73). There was a significant interaction between distance and state (F = 4.5, p = 0.0042), state and time (F = 5.9, p = 0.015), and distance and treatment (F = 3.7, p = 0.012).

Figures 3E and 3F plot the coefficient of variation (COV) for rest and hypercapnic CBF as a function of distance from the injury centre. COV depended significantly on CO_2_ state (F = 21.1, p = 6.3 × 10^−6^) and treatment (F = 7.0, p = 0.012), but not on distance (F = 1.2, p = 0.29) or time (F = 0.28, p = 0.60). There was a significant interaction between state and treatment (F = 11.8, p = 0.00069).

### Vessel density is reduced at 1-day post-TBI and recovers by 4-weeks

Figures 4A and 4B plot the mean vessel length density less than 1 mm from the injury centre, and the vessel density as a function of distance from the injury centre. In Figure 4A, vessel density depended significantly on time (F = 15.4, p = 0.00048) and treatment (F = 28.2, p = 9.6 × 10^−6^), with a significant interaction between time and treatment (F = 9.1, p = 0.0052). Post-hoc group comparisons indicated that vessel density at 1-day post-TBI (410 ± 65 mm/mm^3^) was reduced relative to shams (828 ± 52 mm/mm^3^, p = 1.1 × 10^−5^) and 4 weeks post-TBI (742 ± 45 mm/mm^3^, p = 0.00032). No significant reduction in density was present at 4 weeks post-TBI relative to shams (742 ± 45 mm/mm^3^ vs. 858 ± 30 mm/mm^3^ respectively, p = 0.38).

**Figure 4.**
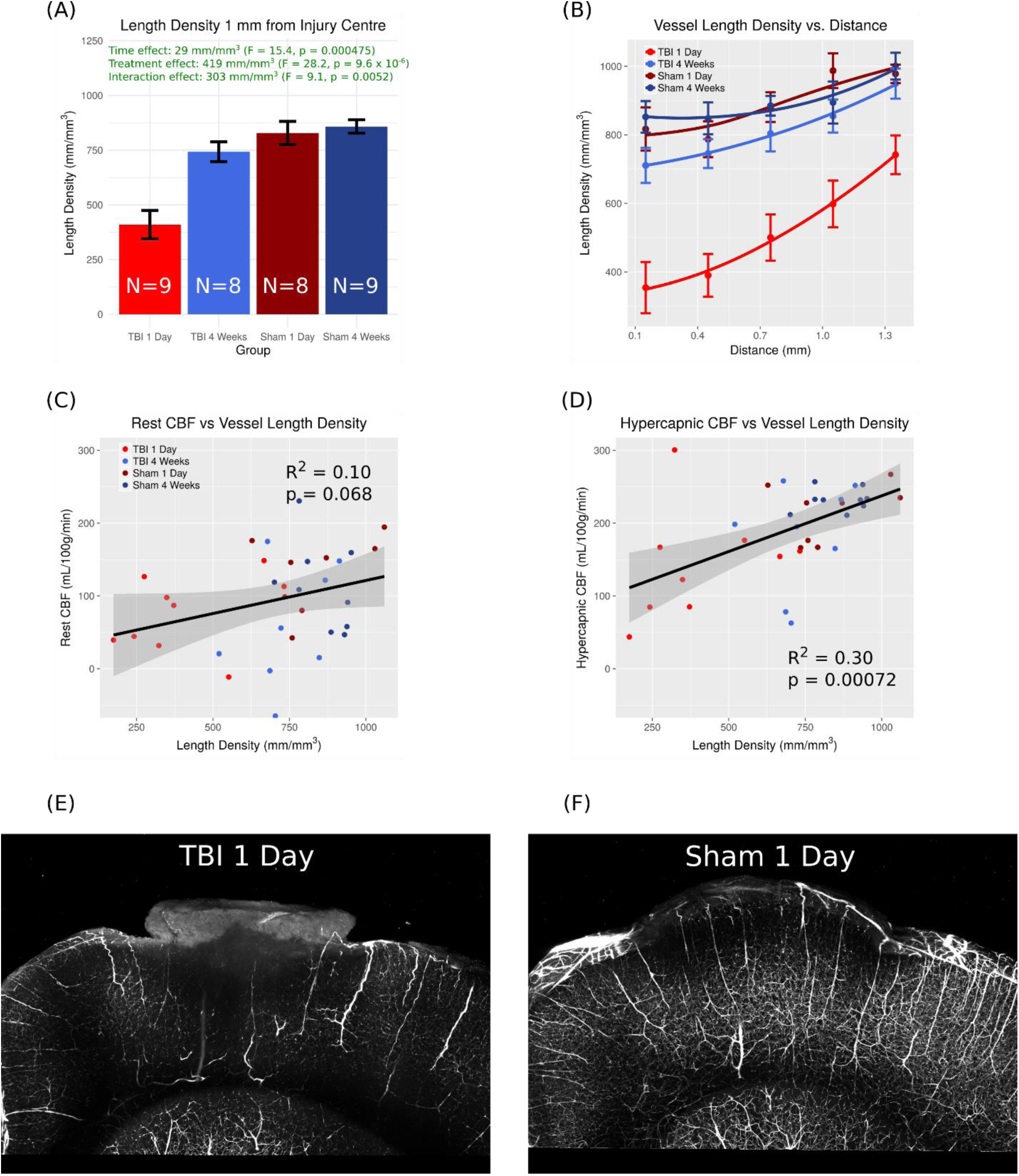
Analysis of Vascular Structural Properties. (A) Mean vessel density (B) Vessel density as a function of distance from the injury centre (C) Rest CBF vs. vessel density (D) Hypercapnic CBF vs. vessel density (E) and (F) MIPs through 400 µm of tissue in a TBI-1-Day and Sham-1-Day mouse respectively. Scale bar = 1 mm. Data points are the mean ± SEM and N refers to the number of mice. Grey bands in C and D represent the 95 % confidence interval.

In Figure 4B, vessel density significantly depended on distance (F = 36.3, p < 2.2 × 10^−16^), time (F = 10.7, p = 0.0027), and treatment (F = 24.8, p = 2.5 × 10^−5^). There was a significant interaction between distance and time (F = 3.0, p = 0.021), distance and treatment (F = 3.7, p = 0.0071), and time and treatment (F = 10.5, p = 0.0029).

Figures 4C and 4D plot the rest and hypercapnic CBF vs. vessel density over all mice. There was a stronger correlation between hypercapnic CBF and density (R^2^ = 0.30, p = 0.00072) compared to rest CBF and density (R^2^ = 0.10, p = 0.068). There was no correlation between rest or hypercapnic CBF and capillary diameter (R^2^ = 0.071, p = 0.13 and R^2^ = 0.0077, p = 0.62 respectively).

Mean capillary diameter over all groups was 4.47 ± 0.05 µm. Mean diameter did not display a time-dependence (F = 2.4, p = 0.13), but did depend on treatment (F = 11.9, p = 0.0017), with the diameter significantly elevated in the TBI condition relative to shams (4.62 ± 0.04 µm TBI vs. 4.33 ± 0.08 µm Shams). There was no significant interaction between time and treatment (F = 0.7, p = 0.41).

Microvascular volume and extravascular distance to the nearest microvessel followed a similar pattern to vessel density (Supplemental Figure 1). Microvascular volume depended significantly on time (F = 21.3, p = 7.0 × 10^−5^) and treatment (F = 9.7, p = 0.0041), with a significant interaction between time and treatment (F = 7.2, p = 0.012). Volume for the TBI-1-Day group (0.74 ± 0.11) was reduced relative to Sham-1-Day (1.29 ± 0.08, p = 0.0016) and TBI-4-Weeks (1.41 ± 0.09, p = 0.00012). There was no statistical difference between TBI-4-Weeks and Sham-4-Weeks (1.41 ± 0.09 vs. 1.45 ± 0.08, p = 0.99). In a plot of volume versus distance (Supplemental Figure 1B), there was a significant dependence on distance (F = 25.8, p = 2.0 × 10^−15^), time (F = 18.6, p = 0.00016), and treatment (F = 6.3, p = 0.018). There was a significant interaction between distance and time (F = 3.8, p = 0.0061), distance and treatment (F = 5.2, p = 0.00073), and time and treatment (F = 7.5, p = 0.010).

The mean extravascular depended significantly on time and treatment (F = 5.2, p = 0.031; F = 18.0, p = 0.00020 respectively), with a significant interaction between time and treatment (F = 7.2, p = 0.012). Extravascular distance at TBI-1-Day (47 ± 5 µm) was elevated relative to Sham-1-Day (26 ± 2 µm, p = 0.00018) and TBI-4-Weeks (33 ± 1 µm, p = 0.012). There was no significant difference between TBI-4-Weeks and Sham-4-Weeks (33 ± 1 µm vs. 28 ± 2 µm, p = 0.69). Extravascular distance depended on distance (F = 16.2, p = 1.3 × 10^−10^), time (F = 5.1, p = 0.031), and treatment (F = 19.0, p = 0.00014) (Supplemental Figure 1D). There was a significant interaction between distance and treatment (F = 5.7, p = 0.00032), time and treatment (F = 7.8, p = 0.0090), and distance : time : treatment (F = 2.6, p = 0.039).

### Correlation of Murray’s Law with blood flow

In Murray’s Law for vessel bifurcations, the cube of the parent vessel radius equals the sum of the cubed radii of the daughters [28, 29, 30]. It assumes vessel networks evolve to minimize the biological work required to transport blood. Many arterial trees closely follow Murray’s Law [31, 32], and deviations from it may potentially indicate abnormal vasculature. Figure 5A is a log-log plot of the average number of vessels greater than or equal to a given diameter, normalized for density, for all vessels within 1 mm of the injury centre. This is termed a cumulative distribution curve. An optimally efficient network with symmetric bifurcations should produce a straight line with a negative slope for each group with a diameter scaling coefficient (absolute value of slope) of 3. For each mouse, a straight line was fit to its log-log plot of vessel diameter distributions, for diameters greater than or equal to 4 µm. 4 µm was selected as the minimum diameter since the average cumulative distributions become horizontal at smaller diameters. The mean diameter scaling coefficients for TBI-1-Day, Sham-1-Day, TBI-4-Weeks, and Sham-4-Weeks were 3.2 ± 0.1, 3.4 ± 0.2, 3.0 ± 0.1, and 3.1 ± 0.1 respectively. When scaling coefficient was modeled as a two-way interaction between time and treatment, there was a trend towards significance for time (F = 3.9, p = 0.058) but no dependence on treatment (F = 1.3, p = 0.27). There was no significant interaction between time and treatment (F = 1.1, p = 0.31). The mean cumulative distribution curve for TBI-1-Day was shifted lower than that of the other groups, indicating a decrease in vessel density (see Figure 5A). Both curves for TBI-1-Day and TBI-4-Weeks showed a ‘dip’ at approximately 8 µm.

**Figure 5.**
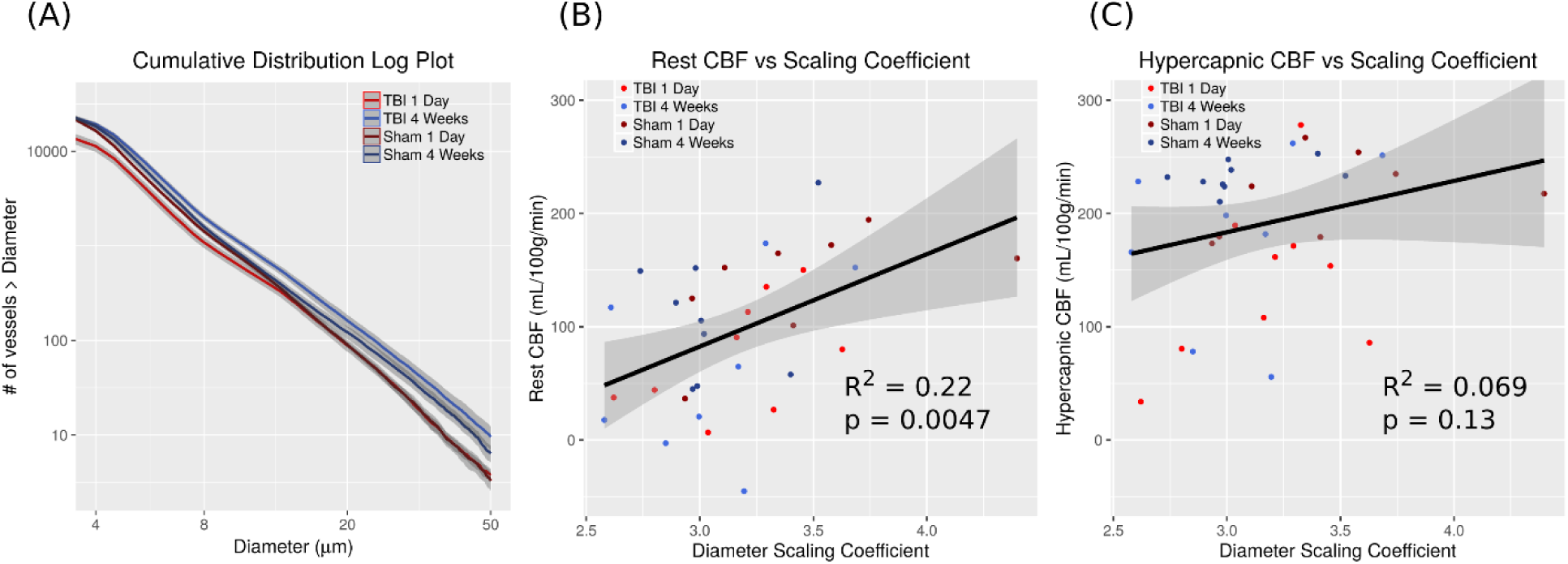
Application of Murray’s Law to imaged vascular networks. (A) Average log-log plots of the cumulative distribution of vessel diameters with standard error. (B) Rest CBF vs. diameter scaling coefficient. (C) Hypercapnic CBF vs. diameter scaling coefficient. Each data point in (B) and (C) represents a single mouse; the grey bands represent the 95 % confidence interval.

Figures 5B and C plot the rest and hypercapnic CBF vs. scaling coefficient for vessels with diameters ranging from 4 – 50 µm. There is a much stronger correlation to the rest compared to hypercapnic CBF (R^2^ = 0.22, p = 0.0047 vs. R^2^ = 0.069 vs. p = 0.13 respectively).

### The vascular architecture possesses a radial pattern at 4 Weeks post-TBI

Figure 6A plots the percentage of vessels as a function of angle (see Methods) within the craniotomy region. There was an emphasis on smaller angles at TBI-4-Weeks relative to the other groups. This is visualized in Figure 6D, with vessels extending radially outwards from the injury centre at the cortical surface in a representative TBI-4-Week mouse, relative to Figure 6C, where there is a more random orientation of vessels in a representative Sham-4-Week mouse. *Radialness* (mean angle) depended significantly on time (F = 23.9, p = 3.2 × 10^−5^) and treatment (F = 33.0, p = 2.9 × 10^−6^), with a significant interaction between time and treatment (F = 18.5, p = 0.00017). There was a dependence of mean angle on group (F = 25.1, p = 2.5 × 10^−8^) (Figure 6B). The mean angle at TBI-4-Weeks (49.4 ± 0.8) was reduced relative to TBI-1-Day (55.4 ± 0.8, p = 1.1 × 10^−6^) and Sham-4-Weeks (55.8 ± 0.4, p = 4 × 10^−7^).

**Figure 6.**
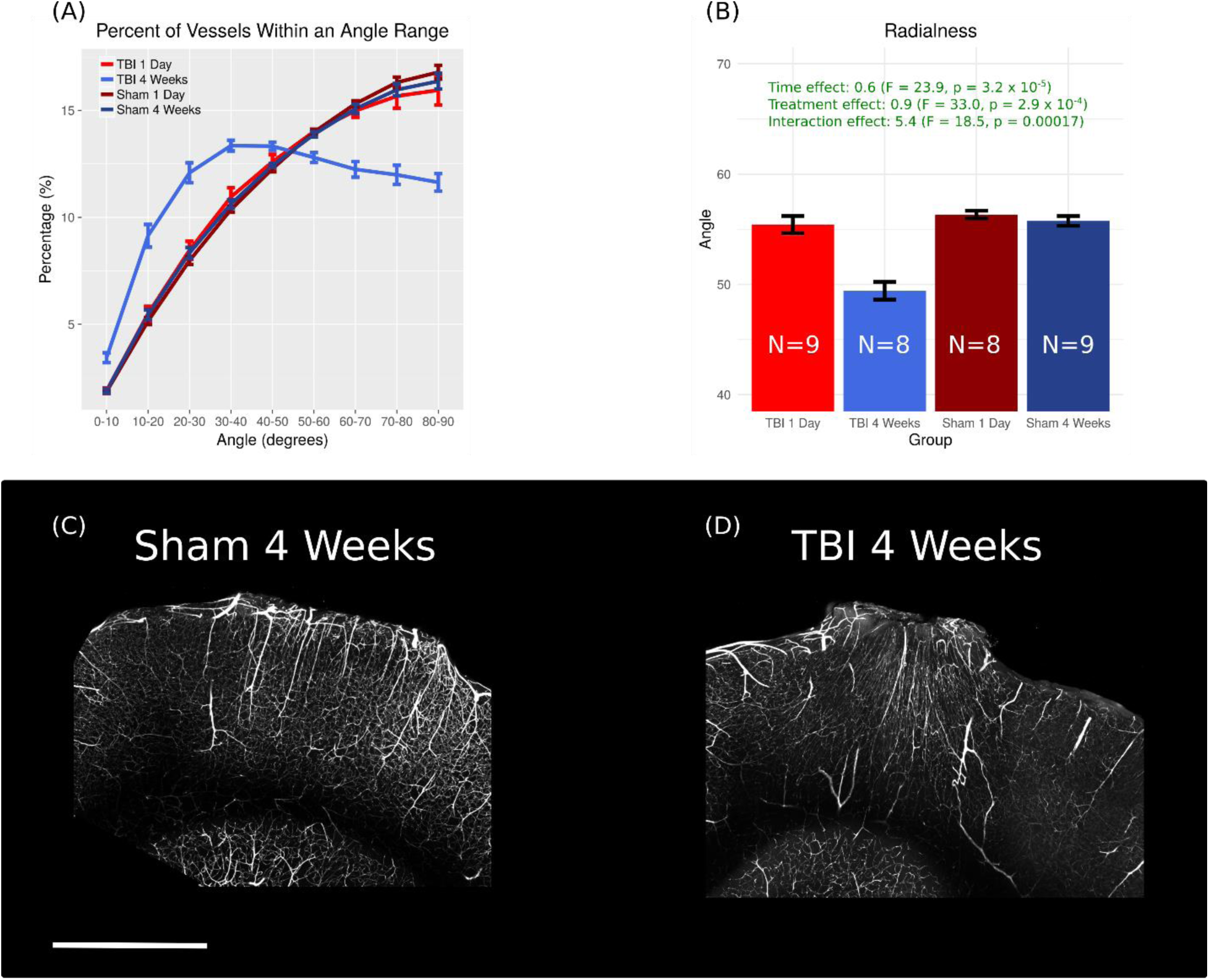
Quantification of the radial vascular architecture. (A) Mean percentage of vessels as a function of angle (B) Mean angle of vessels in injured cortex. (C) MIP through 400 µm of tissue in a Sham-4-Week mouse (D) MIP through 400 µm of tissue in TBI-4-Week mouse. Scale bar = 1 mm. Data points are the mean ± SEM and N refers to the number of mice.

## Discussion

### Biological implications of results

We present a study of vascular architecture and function in a mouse TBI model via a novel methodology for integrating measurements of CBF and microvascular structure. Previous studies of fluid percussion injury in rats described vascular regrowth without concurrent blood flow recovery [5, 9]. Our study is consistent with this result and examines the vasculature in 3D to provide possible explanations for such findings.

While vessel density recovered from TBI by 4 weeks, the vessel network possessed an abnormal radial pattern. It was interesting to note that these vessels, which are likely formed rapidly in the days following injury [33], retained many of the geometric and network properties, such as Murray’s law scaling, that are observed in mature, uninjured vascular networks. Radial patterns of newly formed vessels have been reported in a focal stroke model 6-weeks post-injury [34] and in a mouse CCI model at 7 days [33]. Previous studies have also reported that the size of the injury core reduces with time in CCI [35], which could be due to this radial invasion of new vessels to deliver nutrients to the injured region. The association between this radial pattern and reduced blood flow suggests that this radial patterned network performs more poorly than those found in uninjured animals.

Factors beyond vascular architecture could also influence CBF. Hayward et al. [5] proposed that TBI damages endothelial cells, rendering them incapable of responding to situations requiring hyperemia. TBI also reduces α-SMA in cerebral arteries, which wrap around vessels and contribute to vasomotion and regional CBF flux [36, 37]. The loss of α-SMA is attributed as a potential cause of CBF reductions in trauma [37]. Damaged endothelial cells may poorly conduct blood. In many of the TBI-1-Day mice there were vessels that expressed the endothelial cell fluorescent marker but that the injected FITC gel did not reach (see Supplemental Figure 2). This suggests that these vessels may not be perfused in the resting state in vivo. Nevertheless, gel viscosity does not match blood; there could be increased resistance in the TBI condition leading to a failure of ex vivo perfusion, while in vivo these vessels are still perfused.

In addition to the potential arterial loss of smooth muscle cells, capillaries that lack pericytes often display leakage, dilation, hemorrhage, and reduced flow [38, 39, 40]. In TBI, pericytes migrate away from the vessel wall [41] and are often detached from vessels at least 5 days post-CCI injury [42]. It is possible that a number of these pericytes will remain detached from vessels several weeks later. This would contribute to low blood flow and impaired reactivity.

The positive correlation between rest CBF and diameter scaling coefficient demonstrate that successive branching diameters that decrease more slowly result in a larger CBF, since a larger magnitude coefficient is indicative of diameters decreasing more slowly with successive branching [32]. The poor correlation of scaling coefficient with hypercapnic CBF is possibly due to the fluorescent gel perfusions performed under anaesthetic in the resting state. Although ketamine (anaesthetic used for perfusion) and isoflurane (anaesthetic used for CBF measurements) are both vasodilators [43], hypercapnia under isoflurane exposes mice to an additional vasodilator, CO_2_. In addition to dilating the vasculature, hypercapnia causes a more uniform distribution of vessel diameters and flow [44]. Since the ratio of diameters at a bifurcation drives the scaling coefficient, hypercapnia will affect this ratio, possibly explaining the low correlation. Alterations to 3D vascular structure are further seen in the cumulative distribution plots of Figure 5A, where both TBI groups show an inflection in the cumulative distribution at 8 µm, indicating fewer capillary size vessels. This agrees with the mean increase in diameter calculated in TBI.

Although 3D vascular structure contributes to CBF, vessel density still plays a role. There was a stronger correlation between CBF and vessel density under hypercapnia. This finding could be explained by vessel recruitment whereby the number of capillaries perfused by blood cells increases with stimuli such as increasing metabolic demand or in response to large increases in CBF such as with hypercapnia [45, 46]. Since more vessels are perfused under hypercapnia due to vasodilation, a greater proportion of the vessels imaged with STPT could contribute to CBF in this state and explain the stronger relationship between CBF and vessel density. Previous studies by Hayward et al. [5, 9] correlated only rest CBF with vessel density. At the acute time-point, they found a correlation between density and hippocampal CBF. However, despite an increase of vessel density at 2 weeks, there was a reduction in CBF. This acute correlation is likely due to loss of vessels, while newly formed vessels at later time points may remain un-perfused under resting conditions.

Vascular dysfunction and disruption likely explain the elevated COV, or blood flow variability, measured in TBI. COV increased in TBI close to the injury centre and diminished with distance (Figure 3E, F). Forbes et al. [47] also found increased flow variability at 24 hours in rat CCI. Edema, hemorrhage, and elevated intracranial pressure may compress some vessels, causing redistribution of capillary flows and increased flow heterogeneity [48]. Subarachnoid hemorrhage, a feature of CCI, may cause large vessel vasospasm and concurrent microvessel dilation [49]. We found a 7 % increase in capillary diameter in TBI vs. shams. A combination of redistributed flow, some shrinking vessels, enlarged capillaries, and low flow in regions of directly damaged vasculature and hemorrhage may contribute to the CBF heterogeneity observed.

Despite vessel loss at 1 day, our findings imply that tissue remains partially viable post-injury. Park et al. [4] showed that if an injury is sufficiently severe, the vasculature will not regrow weeks after TBI. The extravascular distance to the nearest microvessel was elevated 1-day post-TBI relative to other groups in our experiments. The mean extravascular distance in our study at 1 day was 47 ± 5 µm, while normal synaptic structure is maintained if a dendrite is within 80 µm of a microvessel [50]. Many dendrites would be able to receive oxygen and nutrients from the vascular network since they are within the 80 µm limit. This would enable the tissue to regrow and repair to some extent, which was found in the TBI-4-Week mice. Tissue recovery will also partially depend on flow through the network, which remains reduced at 4-weeks post-TBI. Further experiments are needed to determine the status of dendrite and neuron function. However, recovery of vessel density and the absence of a necrotic cavity as per more severe CCI models [51] suggests the presence of at least partially viable tissue.

### Technical notes

As the methodology used for vessel density measurements is novel, it is interesting to compare these observations with literature reports. Mean vessel length density at Sham-4-Weeks (858 ± 30 mm^3^) reported here is comparable to that obtained in Tsai et al. [22] from sucrose-cleared tissue slabs (880 mm/mm^3^), Boero et al. [52] from thin sections (700 – 1200 mm/mm^3^), and Lugo-Hernandez et al. [21] from whole brain microvascular reconstructions (922 mm/mm^3^). Tsai et al. [22] report smaller capillary diameters at 3.5 – 4.0 µm, compared to a mean of 4.5 µm, despite usage of identical perfusion materials. This could be due to their quantifying the lumen diameter whereas the diameter measurement reported here include the vessel wall. These diameter differences likely explain the greater microvascular volume (1.45 % at Sham-4-Weeks vs. 1 % in Tsai et al).

There are drawbacks to STPT. As an ex vivo technique, it necessitates design of a cross-sectional study. Unlike optical clearing where the tissue may be sectioned for histology following imaging, subsequent histological analysis is not possible with STPT. This prevents examination of relevant molecular and cellular-level processes beyond vasculature structure, such as presence of pericytes or smooth muscle cells, which may impact blood flow.

Finally, while this study was conducted on both male and female mice, it was underpowered to examine sex differences. It was recently reported in a mouse CCI model that vessel network complexity and vessel numbers differed by sex post-injury [8]. No sex dependence was found in the present study: there was no statistical dependence of CBF on sex; both sexes demonstrated vascular loss at 1 day, followed by recovery to sham levels; and both sexes exhibited a radial vascular pattern at 4 weeks (data not shown).

## Conclusions

Traumatic brain injury leads to widespread changes in the cerebral vascular system, whose functional and structural characteristics evolve over the days and weeks following injury. In the present study, by examining the functional characteristics of the cerebral circulation using non-invasive MRI measurements of cerebral blood flow and ex vivo measurements of the 3D structure of the blood vessels networks proximal to the injury, we found that these two aspects of the circulation were related in more complex ways than can be summarized as a simple magnitude of injury. Remodeled vessel networks following injury demonstrated a highly regular hierarchical structure that is characteristic of healthy vessel networks but did not recover the level of vascular function seen in sham treated animals. By other morphological measures, however, these remodeled networks were highly abnormal and possessed a characteristic radial network pattern. In developing future therapies to promote recovery of vascular function following injury, enhancing vessel growth may not be sufficient as the 3D architecture is an important factor in the overall performance of the remodeled network.

## Author Contribution Statement

All authors were involved in the design of experiments. JS and LSC conducted the experiments. JS, LSC, and JGS analyzed the data. JS was the primary manuscript author. LSC, MMK, BS, and JGS reviewed and edited the manuscript.

## Supplementary Information

**Supplemental Table 1.**
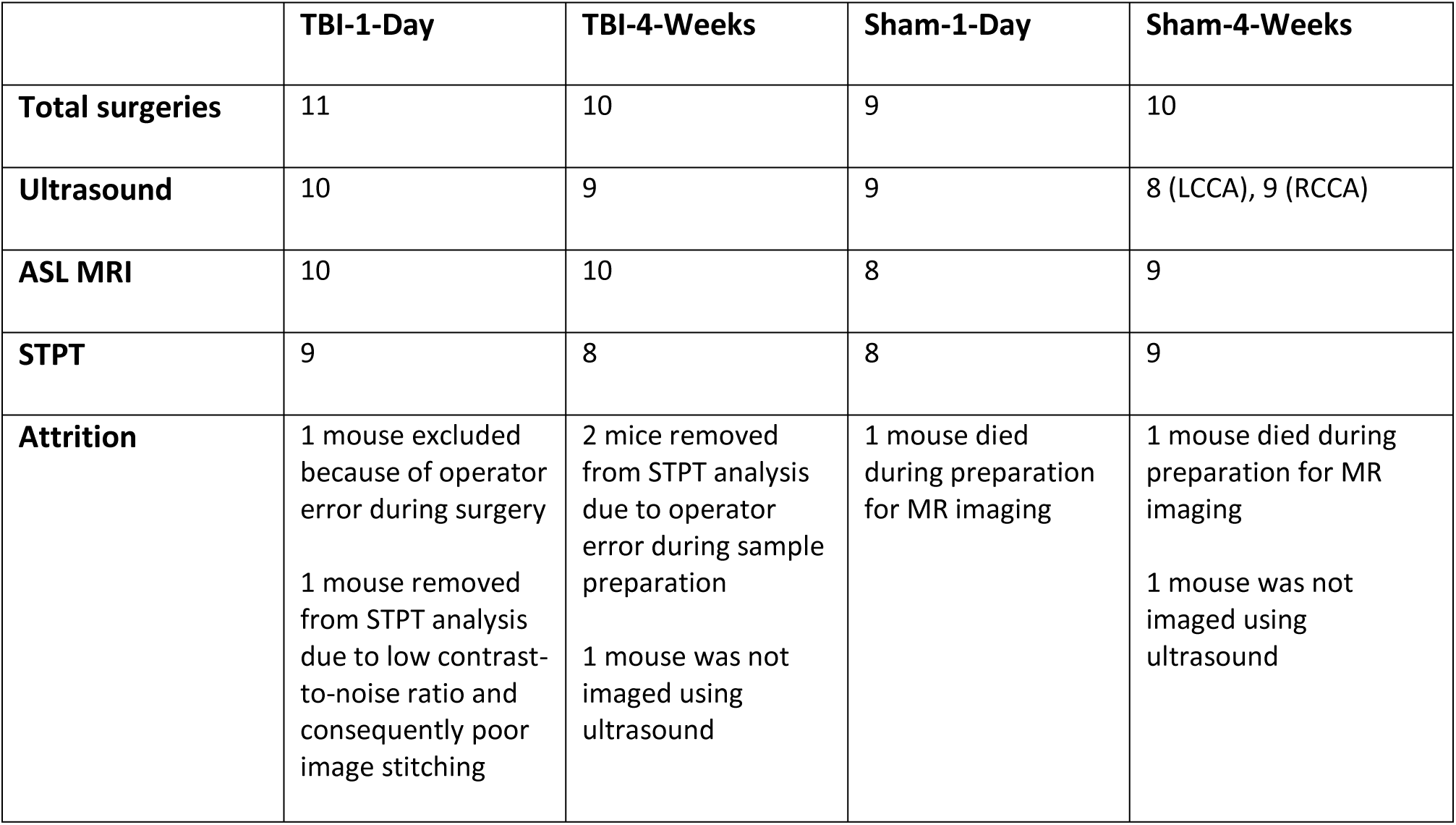
Number of mice assigned to each group (TBI-1-Day, TBI-4-Weeks, Sham-1-Day, Sham-4-Weeks), along with numbers lost to attrition. A total of 40 mice were used for the study.

|  | **TBI-1-Day** | **TBI-4-Weeks** | **Sham-1-Day** | **Sham-4-Weeks** |
| --- | --- | --- | --- | --- |
| **Total surgeries** | 11 | 10 | 9 | 10 |
| **Ultrasound** | 10 | 9 | 9 | 8 (LCCA), 9 (RCCA) |
| **ASL MRI** | 10 | 10 | 8 | 9 |
| **STPT** | 9 | 8 | 8 | 9 |
| **Attrition** | 1 mouse excluded because of operator error during surgery<br>1 mouse removed from STPT analysis due to low contrast-to-noise ratio and consequently poor image stitching | 2 mice removed from STPT analysis due to operator error during sample preparation<br>1 mouse was not imaged using ultrasound | 1 mouse died during preparation for MR imaging | 1 mouse died during preparation for MR imaging<br>1 mouse was not imaged using ultrasound |

**Supplemental Figure 1.**
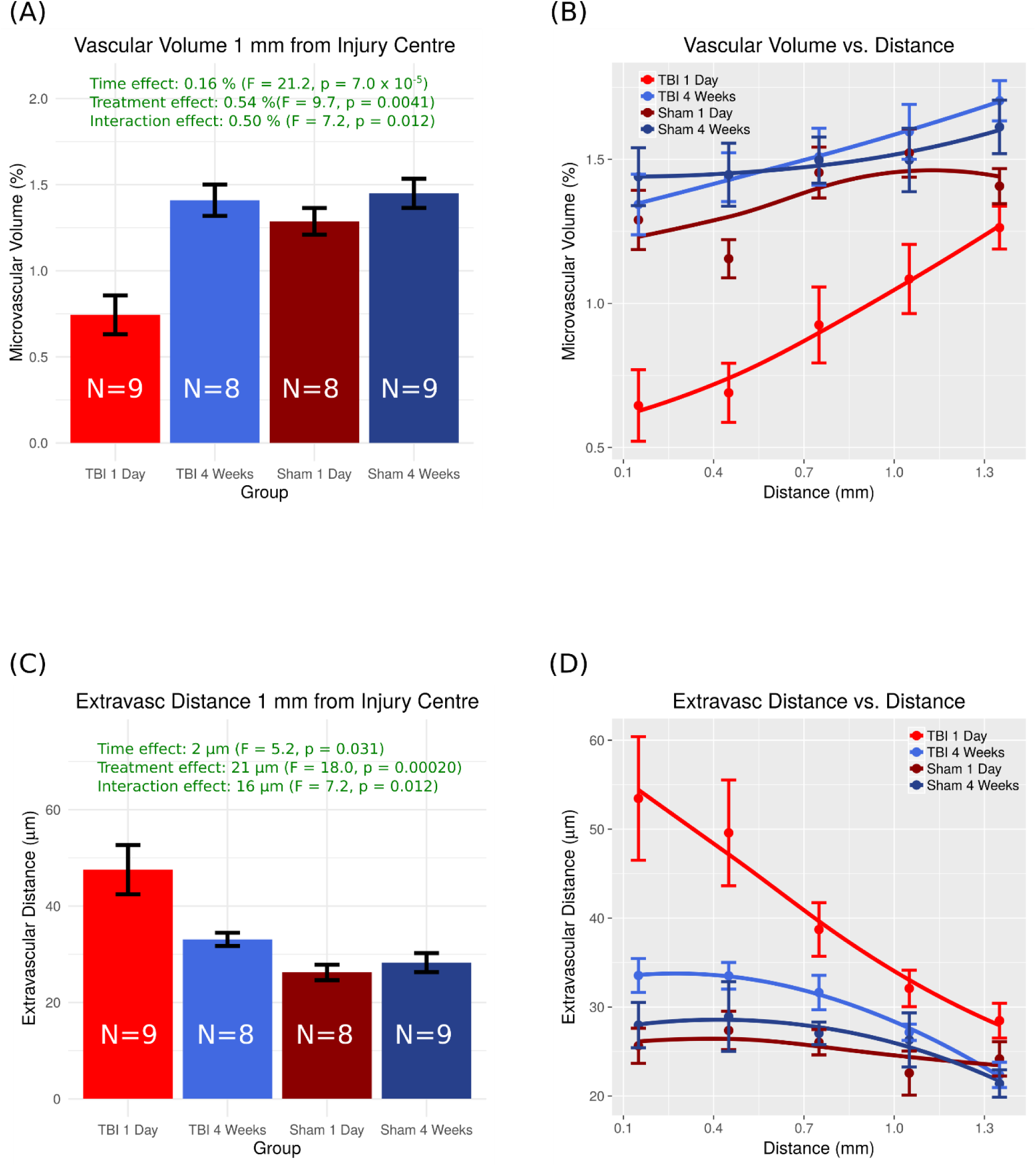
(A) Mean vascular volume. Volume was reduced at TBI-1-Day vs. shams and TBI-4-Weeks (B) Vascular volume as a function of distance from the injury centre (C) Mean extravascular distance to the nearest microvessel. (D) Extravascular distance to the nearest microvessel as a function of distance from the injury centre.

**Supplemental Figure 2.**
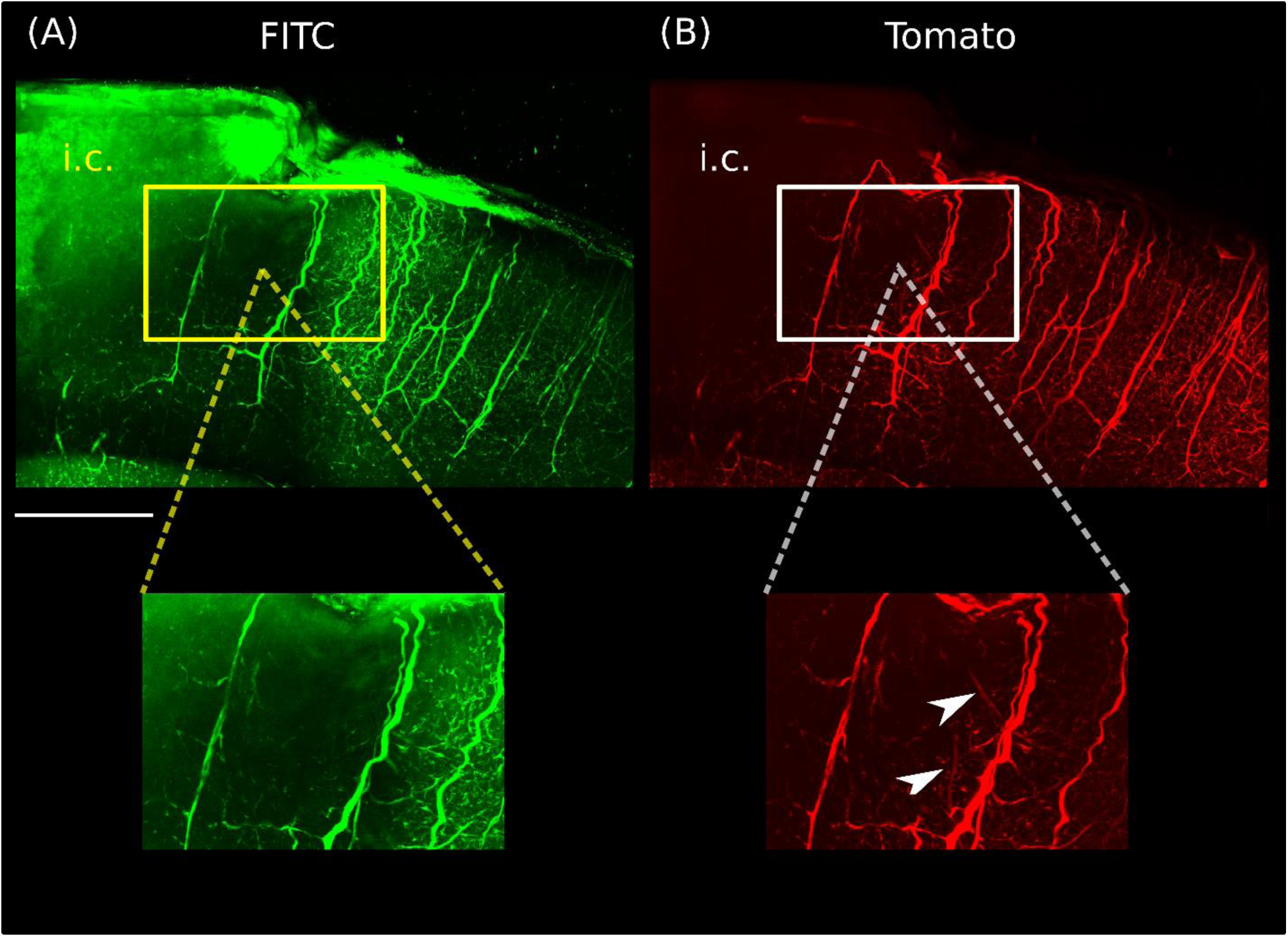
(A) MIP (700 µm tissue thickness) through STPT data for a TBI-1-Day mouse showing the FITC fluorescence from the perfused gelatin detected following spectral demixing. (B) MIP of the same tissue region as (A) demonstrating tomato fluorescence. i.c. represents the injury core region. The boxes in (A) and (B) show the vasculature in the penumbra region surrounding the core. While a number of tomato-fluorescent vessels are perfused within the penumbra, others are not successfully perfused (see white arrows in inset from tomato image).

